# Evaluating Field Corn Yield and Plant and Soil Nutrient Concentrations Under Application of Synthetic Fertilizer and Dairy Manure

**DOI:** 10.1101/2025.09.28.676050

**Authors:** Tajamul Hussain, Muhammad Fraz Ali

**Affiliations:** Oregon State University, United States; Northwest A&F University, China

**Keywords:** Field corn, manure, yield, nutrient recovery, leaching, nitrate-N

## Abstract

Application of manure in field corn has the potential to sustain corn yields and reduce nutrient leaching in soil profile. A field trial with randomized complete block design was conducted on Adkins fine sandy loam soil to evaluate the impact of application of manure and synthetic fertilizer on nutrient concentrations (N, P, K and S) in plant and soil and field corn yield. Experimental treatments included an application of synthetic fertilizer (NPK) and dairy manure application at 5-, 10- and 15-tons acre^-1^ in addition to a non-fertilized control. All the manure was applied before planting. Corn was manually harvested, and plants were separated into leaves, stems and cobs to determine dry weights. Post harvest soil sampling was performed at 0-30, 30-60 and 60-90 cm soil depths. Results indicated that in-season leaf nutrient concentration was significantly different among applied treatments. Application of synthetic fertilizer resulted in the highest plant height (116 in) and produced higher corn yield (45.5 tons acre^-1^) compared to control and application of dairy manure. Application of manure at 5 tons acre^-1^ produced higher corn yield (35.5 tons acre^-1^) compared to manure application at 10 (25.9 tons acre^-1^) and 15 tons acre^-1^ (26.1 tons acre^-1^). A similar trend was observed for leaf, stem and cobs fresh and dry weights. Nutrient recovery was higher under application of synthetic fertilizer followed by application of manure at 5 tons acre^-1^. Soil nutrient analysis indicated no significant impact on N, P, K and S concentration among treatments except for NH_4_^+^–N. However, nutrient concentration significantly varied under different soil depths. Results suggest that a combination of synthetic fertilizer application and manure might be a practical approach for balanced nutrient supply for field corn. Further investigations are necessary to explore the potential of manure application to ensure balanced nutrient supply, improved yields and reduced nutrient losses in field corn.

**Graphical Abstract:** 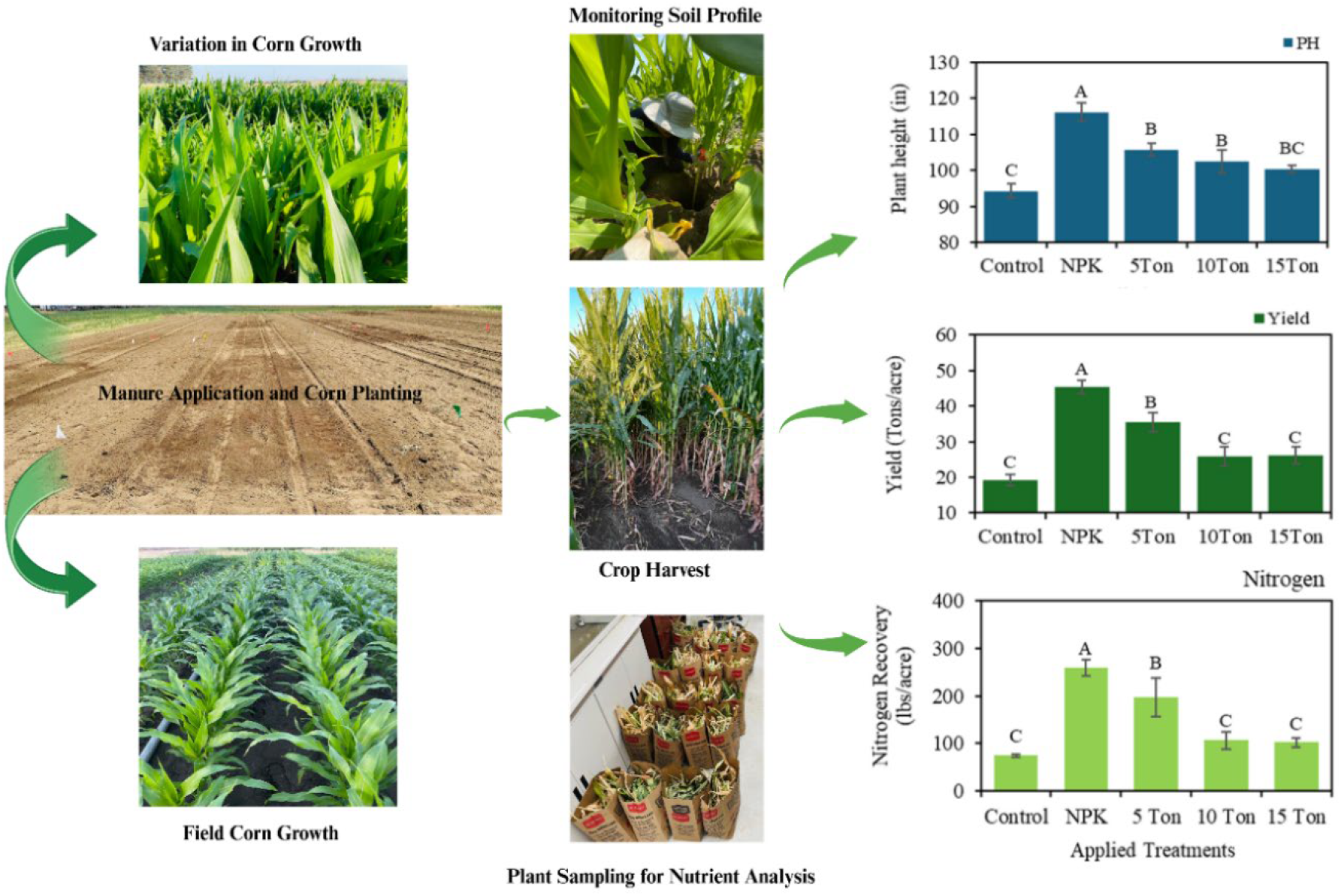

***Credit:*** *Photographs taken and experimental data generated by T. Hussain during postdoctoral research at OSU*.

## 1. Introduction

Corn silage is an important component of animal feed in dairy industry across United States including Oregon. Approximately 40% of total domestic corn is used in animal feed (USDA, 2025) and statistics indicate that silage corn was harvested from 6.1 million hectares of land (Villa, 2025). Silage corn is considered as high energy feed for dairy cattle due to its dry matter content and palatability (Sullivan et al., 2023). Research has demonstrated that feeding corn silage to dairy cows helps in improving feed intake, milk production, and milk protein content (Kok et al., 2024; Tharangani et al., 2021; Bach et al., 2021). Efficient production of silage corn is linked to effective nutrient management to meet crop demands. However, the production potential of Columbia Basin region is challenged by its coarse-textured soils, which exhibit low soil-water and nutrient holding capacity and low soil organic matter. Synthetic fertilizers are the main source of nutrients for silage corn production (Baghdadi et al., 2018). To compensate for inherent soil limitations and achieve optimal yields, growers often apply large quantities of synthetic fertilizers. However, the applications of these synthetic fertilizers may lead to soil degradation and nutrient leaching, posing a serious threat to agricultural productivity and sustainability (Hina, 2024; Tufail et al., 2024; Singh and Craswell, 2021).

Achieving and sustaining optimal corn yields depends heavily on effective crop nutrient management that aligns soil nutrient availability with crop uptake patterns. Corn has a high demand for essential macronutrients like nitrogen (N), phosphorus (P) and potassium (K), which are critical for supporting key physiological processes and optimal plant growth and development (Cao et al., 2025; Singh et al., 2023; Cui and Tcherkez, 2021). Specifically, N is vital for leaf development and chlorophyll synthesis (Leghari et al., 2016), P promotes root growth (Liu, 2021), and K regulates water utilization (Xu et al., 2021; Kandil et al., 2020). Corn silage has greater potential for nutrient recovery, and it can remove 205, 32 and 192 lb N, P and K per acre, respectively (Sullivan et al., 2023). Plant growth and yield potential can be significantly restricted by deficiencies or imbalances of these essential nutrients. To meet these intensive nutrient demands, growers have historically relied on synthetic, inorganic and organic fertilizers (Wang et al., 2024; O’Brien and Hatfield, 2019; Ciampitti and Vyn, 2014). However, the overuse of synthetic fertilizers to achieve the maximize yields undermines the parallel goal of preserving long-term soil fertility (Mustafa et al., 2023). The dependency on synthetic fertilizers also presents significant economic and environmental challenges, including nutrient leaching that contaminates groundwater and runoff that contributes to eutrophication of surface waters. Therefore, nutrient management strategies must not only ensure the timely availability of nutrients and improved crop yields but also promote soil health.

Manure management is also a substantial issue as managing the large volumes of manure generated by dairy farmers poses both economic and environmental challenges. The immense quantities of manure generated represent a valuable, renewable source of plant nutrients and organic matter (Key et al., 2023; Park et al., 2019). In recent years, dairy farm manure has drawn significant attention in an effort of sustainable agriculture, with potential to improve soil health, enhance crop yields and mitigate environmental impacts (Key et al., 2023; Verma et al., 2019). By recycling nutrients within the agricultural system, dairy manure application can reduce the dependence on synthetic fertilizers and transform a waste product into a resource, creating a more circular and sustainable nutrient economy. Manure is valuable nutrient resource that helps to build soil organic matter, improve soil health and enhance nutrient cycling when applied at right time and rate (Ramos Tanchez, 2023). Manure improves soil fertility and enhances microbial activity on short them which leads to improved soil structure. Whereas in the long term it supplies nitrate and ammonium that facilitates crop productivity (Dungan et al., 2024; Wang et al., 2023, Khairunisa et al., 2023). Despite these potential benefits and research efforts on fertilizer and manure application in corn, a critical knowledge gap persists. There is a lack of comprehensive field scale studies in the Columbia Basin that directly explain efficiency of dairy manure compared to synthetic fertilizers by evaluating corn yield, nutrient recovery, soil nutrient dynamics and potential environmental impacts. Columbia Basin counties have a rich manure source, e.g., from dairy farms. The cost of the manure is meager (or no cost) as these farms are suffering environmental regulations on disposing of the manure. Applying manures to agricultural fields will be a win-win strategy to help these farms lessen the environmental pressure and benefit field crop growers to improve soil quality. The use of manure in fields represents an opportunity to recycle an existing by-product capable of providing agronomic and environmental benefits. Although most manure typically contains lower concentrations of N, P, and K than commercial fertilizers, it can be a valuable source of organic matter, and other essential macro-and micro-nutrients (Timsina, 2018). Short-term or long-term effects of manure application on key soil properties in this Columbia Basin remain unclear due to the limitation of brief study periods and limited experimental factors. Therefore, research evidence is needed to quantify whether dairy manure can be a viable substitute for synthetic fertilizers in achieving optimal corn yield while enhancing soil fertility and mitigating environmental risks in the Columbia Basin’s unique agroecosystem.

There is limited information available on how application of dairy manure affects corn growth, nutrient uptake and nutrient dynamics in sandy soils. While manure provides valuable nutrients, a particular concern is that the gradual nutrient release from manure might not be able to meet the crop requirement during the peak growing period. In addition, impact of manure application on soil nutrient dynamics remains unexamined in the Columbia Basin. To fill this critical knowledge gap, this study aimed to provide a field-based evaluation of manure management strategies for silage corn. Therefore, the objectives of this research were 1) to assess the influence of synthetic fertilizer and dairy manure application on corn growth, yield and nutrient recovery, 2) to determine the effects of these nutrient sources on soil nutrient dynamics and mobility in soil profile and 3) to identify optimal dairy manure application that can sustain high yield while reducing environmental risks. The current research will provide critical insights for developing effective and sustainable manure and nutrient management for silage corn production in Columbia Basin and similar regions, with implications for optimizing productivity, improving soil health and preserving environment.

## 2 Material and Methods

### 2.1 Field Experiments

Field trial was conducted at the field research area of Hermiston Agriculture Research and Extension Center, Oregon State University (latitude: 45° 49’ 1.63” N, longitude: 119° 16’ 51.24” W) during 2024 growing season. The soil of the experimental field is Adkin fine sandy loam. The experiment was conducted in a randomized complete block design with four replications having a total of 20 plots. The field was prepared using disc plough, disc harrow and rotary tiller twice. Experimental treatments included 1) a control (without fertilizer), 2) chemical fertilizer application (NPK), 3) dairy manure application at 5-ton acre^-1^, 4) dairy manure application at 10-ton acre^-1^, 5) dairy manure application at 15-ton acre^-1^. Each experimental treatment was designated in a 25 × 8 ft plot having four rows of corn having 2.5 ft of row-row distance surrounded by buffer area. Planting was performed on May 23, 2024, and corn was planted with a two-row mechanical planter at a plant-to-plant distance of 7 inches. All manure was applied after field preparation and before crop planting in designated plots. Whereas chemical fertilizer application was performed during the season. Irrigation was applied using manually operated sprinkler irrigation system and the irrigation amount was based on the estimated evapotranspiration data provided by the local meteorological station (HRMO) at the OSU-HAREC (AgriMet: https://www.usbr.gov/pn/agrimet/graphs.html). Weather data was also obtained from Agrimet. Mean minimum and maximum air temperature and relative humidity during the crop growth season ranged 5.3-24.3 and 19.7-41.3 °C and 28-68 %, respectively. The experiment received 588 mm of irrigation water with only 8.1 mm of rainfall. Standard practices were employed to control insects and pests.

### 2.2 Data Collection

During the crop growth season, the crop growth was monitored by taking four leaf samples from each plot. The plant samples were taken to evaluate the effect of various treatments on nutrient concentration. The crop was manually harvested during early September when the average soil moisture of corn reached 65%. Plant height was measured from the base of the stem to the tip of the flag leaf. Plants in the 4 ft length from central two rows were harvested to determine total aboveground biomass whereas three hills having two stalks per hill were selected to determine fresh and dry weights of plant parts including leaf, stems and cobs (Figure 1). Plant samples were dried in oven at 65 °C to achieve a constant weight. Dried samples were then stored for laboratory analysis to determine the nutrient concentrations and recovery. Soil sampling was performed in all experimental plots to determine soil nutrient concentrations in soil. Samples were obtained from the harvested area at 0-30, 30-60 and 60-90 cm soil depths in each plot.

**Figure 1:**
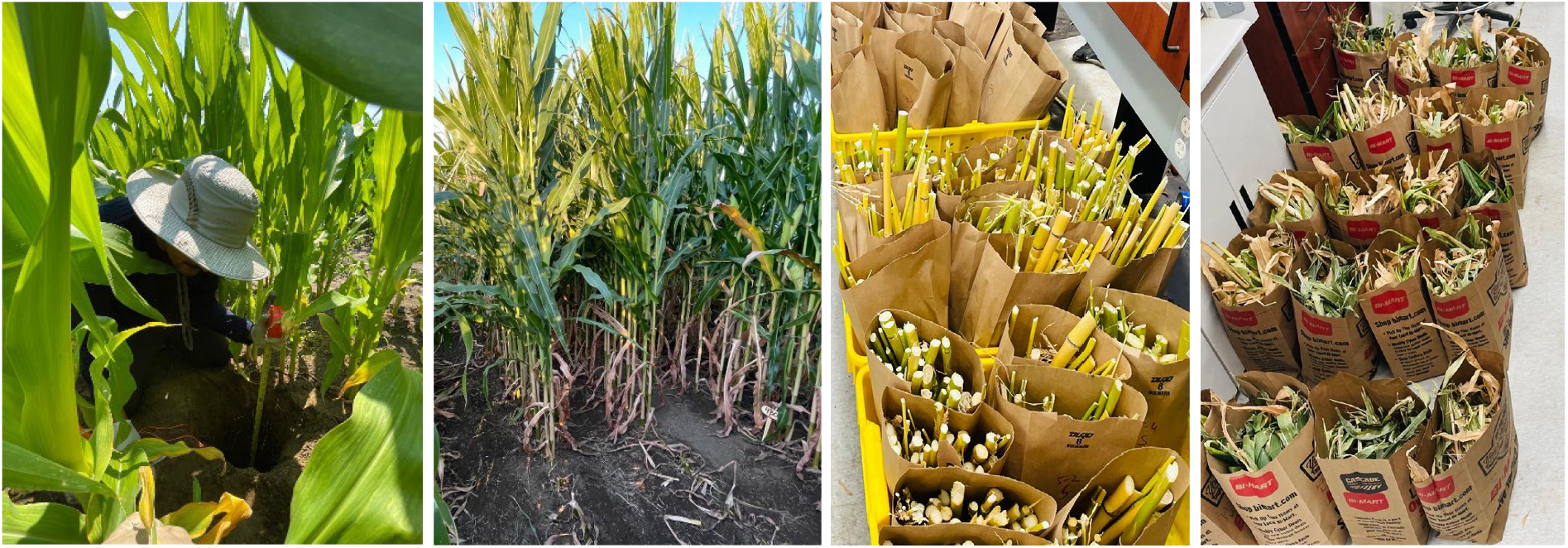
Monitoring soil profile, plant sampling at harvest and sample preparation for nutrient analsis. (Credit: *Tajmul Hussain*)

### 2.3 Nutrient Concentration and Analysis

Dried plant samples including leaf, stem and cobs were ground to pass a 2-mm sieve. Soil was cleaned, samples were prepared and passed through a 2-mm sieve. NO_3_^−^–N, NH_4_^+^–N, P_2_O_5_, K^+^ and SO_4_–S in samples were determined following the NAPT S-3.10, NAPT S-4.10, NAPT S-5.11 and NAPT S-6.12 standard analytical procedures for testing (Miller et al., 2013).

### 2.4 Statistics

Statistical analysis was performed using Statistix 8.1 software to evaluate the effects of various applied treatments on field corn growth, yield and plant and soil nutrients through conducting analysis of variance and mean comparisons using the least significant difference test. Means were considered significantly different at *p* values < 0.05.

## 3. Results and Discussion

### 3.1 In-seasons Leaf Nutrient Concentration

In-seasons leaf N and K (Figure 2A) and S (Figure 2B) concentration were significantly different whereas no significant difference was observed for P concentration under applied treatments. Leaf N, K and S concentrations were highest in NPK treatment. This indicates that application of synthetic fertilizer directly influenced N, K and S in plants. Singh et al. (2023) found that nutrient concentration in corn influenced at different crop stages. Leaf N concentration was higher in manure application at 5 Tons whereas it was statistically similar under 10 ton, 15-ton manure application and in control treatment. These findings suggest that application of 5-ton manure application was sufficient to meet the crop in season N, P, K and S demand and higher manure application may lead to higher nutrient in soil profile that may be prone to leaching if not used by the crop in remaining season. Leaf K concentration was statistically similar in all manure applications and control treatments. This suggests with the experimental condition and native soil K supply or K availability from manure treatments did not significantly influence leaf K. This is also possible due to soil buffering capacity to maintain consistent levels or supply across different treatments. Leaf S concentration indicated a similar trend to leaf N concentration and higher manure applications resulted in similar results with control treatment (Figure 2B). Higher manure application rate enhances soil organic matter and microbial activities facilitating mineralization and sulfur availability. A similar trend with N, S concentration indicated possible linkage in silage corn. This also suggests that application of 5-ton manure was sufficient to supply maximum sulfur for field corn and sulfur contribution from higher manure applications may have limited impact on leaf S beyond certain thresholds.

**Figure 2:**
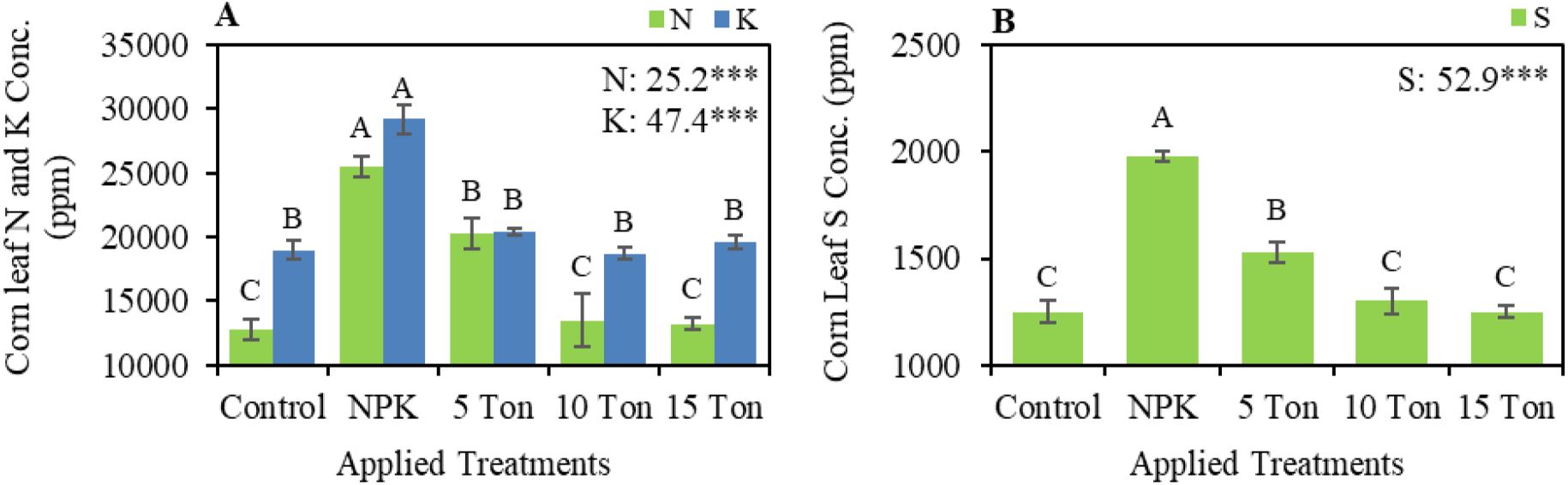
Impact of synthetic fertilizer and manure application rates on in-season leaf nutrient concentrations including leaf N and leaf K (**A**) and leaf S (**B**). F-values are given in the charts with significance. Vertical bars indicate ± standard errors for means of four repetitions. Letters above the bars indicate the significant (*p* < 0.05) differences among applied treatments.

### 3.2 Corn Growth and Yield

Statistical evaluation indicated that corn growth and yield parameters including plant height, leaf, stem and cobs fresh and dry weights were significantly different under the effect of applied treatments Table 1. The highest plant height was observed under NPK treatment that was 23% higher than that of control. Plant height under manure application was statistically similar and lower than the NPK application, however it was higher than control by 6, 9 and 12% under 15 tons, 10 tons and 5 tons manure application, respectively (Figure 3A). Higher plant height in NPK treatment followed by application of manure at 5-ton indicates that a balanced supply of readily available nutrients is critical for improved growth and application of manure may not adequately supply required nutrients due to complex interactions and processes in soils. Therefore, combination of synthetic fertilizers and manure applications may improve nutrient availability and ensure optimal plant growth. Syamsiyan et al. (2024) suggested that application of manure with inorganic fertilizer improved growth and yield of sweet corn. Similarly, Baghdadi et al. (2018) who tested chicken manure and chemical fertilizers suggested practicing integrated application of manure and chemical fertilizers to achieve optimal yield and quality of corn silage.

**Table 1.** *F*–values and significance obtained through analysis of variance of corn growth and yield parameters including plant height, silage yield, leaf fresh weight (LFW), leaf dry weight (LDW), stem fresh weight (SFW), stems dry weight (SDW), cobs fresh weight (CFW) and cobs dry weight (CDW).

|  | PH | Yield | LFW | LDW | SFW | SDW | CFW | CDW |
| --- | --- | --- | --- | --- | --- | --- | --- | --- |
| Treatment | 264.3*** | 417.8*** | 12.3** | 0.8** | 20.4** | 0.8** | 63.9*** | 9.8*** |
| Error | 21.4 | 23.8 | 1.4 | 0.1 | 2.2 | 0.1 | 4.1 | 0.6 |
| CV | 4.5 | 16.0 | 24.8 | 13.3 | 14.0 | 13.7 | 20.6 | 23.1 |

\*\*\*: $p < 0.001$ , \*\*: $p < 0.01$ , \*: $p < 0.05$ , ns: non-significant difference at $p \geq 0.05$ .

**Figure 3:**
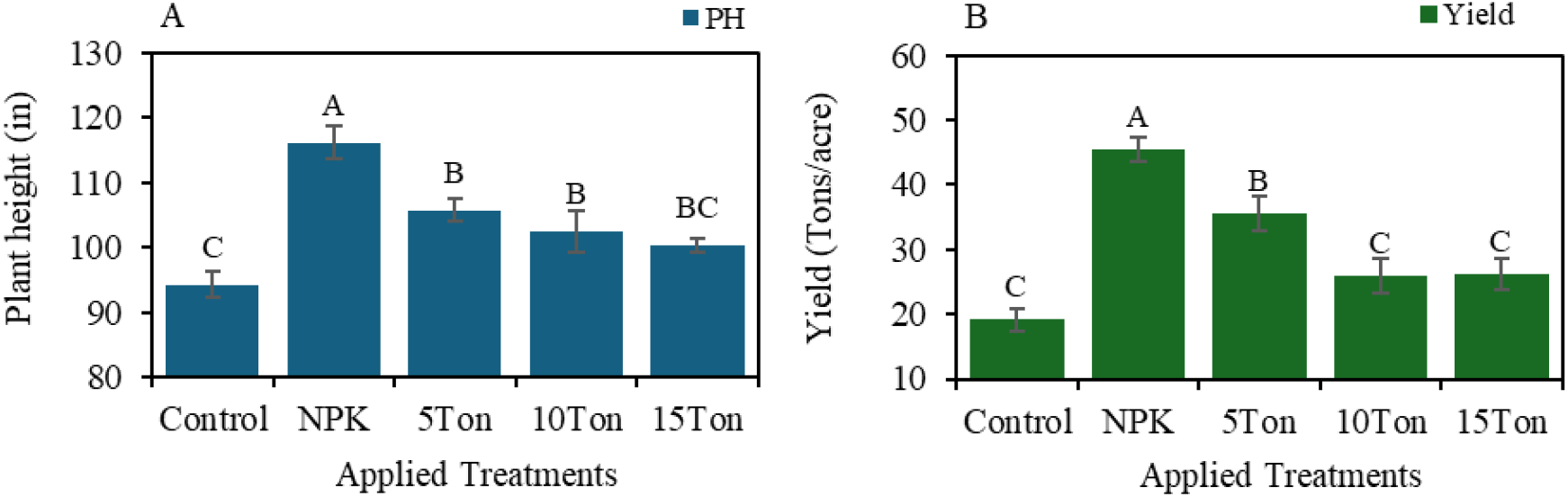
Impact of synthetic fertilizer and manure application rates on plant height (A) and yield (B) of field corn. Vertical bars indicate ± standard errors for means of four repetitions. Letters above the bars indicate the significant (*p* < 0.05) differences among applied treatments.

Corn yield was also indicated a similar trend with respect to NPK application and highest yield 45.5 tons acre ^-1^ was observed under NPK treatment. Corn yield under NPK treatment was 137% higher compared to control and it under manure applications was higher by 36, 35 and 85% under 15 tons, 10 tons and 5 tons manure application respectively (Figure 3B). Under manure application maximum corn yield was achieved at 5 tons manure application whereas yield was statistically similar under manure application at 10- and 15-tons manure application and control (Figure 3B). This trend indicated that application of NPK met the corn nutrient requirements whereas application of manure alone did not sufficiently provide required nutrition and application of manure beyond 5 tons possibly exacerbated negative impacts possibly due to increased nutrient imbalance or soil microbial activity. This might also be attributed to field conditions and soil properties. Previous research conducted by Ramoz

Tanchez et al. (2023) indicated that when the field had sufficient nutrient supply, corn silage did not respond to manure application. A suitable manure application rate combined with reduced synthetic fertilizer application may be strategic approach to enhance production efficiency of silage corn. A recent study conducted by Wang et al. (2025) suggested that the yield and quality of maize improved under combined application of organic fertilizers and chemical nitrogen. Similarly, Qu et al. (2025) concluded their recommendations stating that although manure application has potential to enhance yields, it indicated poor yield stability whereas integrated manure application with mineral fertilizers enhanced yield stability and soil health.

Leaf and stem fresh weight and dry weights indicated a similar trend, and maximum weights were observed under NPK application that were 134%, 63% and 80% and 68% higher than the control for leaf and stem fresh and dry weights respectively (Figure 3A,B). Under manure application, leaf and stem fresh as well as dry weights were higher under 5 tons manure application followed by 15 tons and 10 tons. In comparison to control, leaf and stem, fresh and dry weights were 54%, and 31% and 37% and 33% higher at 5 tons manure application, respectively whereas they were 23%, and 23% and 35% and 28% higher at 15 tons manure application. Leaf and stem, fresh and dry weights were decreased under 10% manure application, however, they were still higher than that of control by 13%, and 14% and 24% and 26%, respectively (Figure 4A,B). Cobs were also separated, and the results indicated a similar trend with leaf and stem fresh and dry weights. The highest cob fresh and dry weights were observed under NPK application that were 179% and 234% higher compared to control. Under manure application, 5 tons of manure resulted in higher cobs fresh and dry weights that were 145% and 197% higher than control. Cobs fresh and dry weights were decreased under 10- and 15-tons manure applications however, they were higher by 60% and 84% and 39% and 60% for 10- and 15-tons manure applications compared to control (Figure 4A,B). A similar trend in fresh and dry weights of plant parts with corn yield indicated a similar corn response to applied treatments. Therefore, it is critical to consider optimal manure application for balanced nutrient availability and optimal yields.

**Figure 4:**
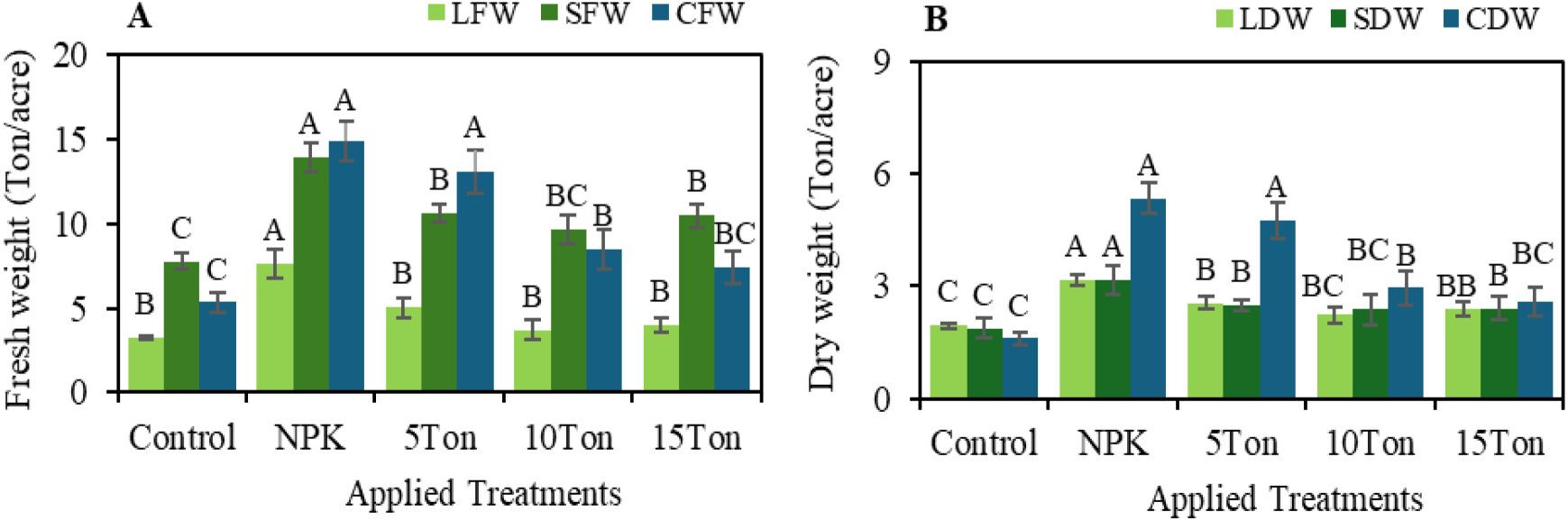
Impact of synthetic fertilizer and manure application rates on leaf, stem and cobs fresh (A) and dry (B) weights of field corn. Vertical bars indicate ± standard errors for means of four repetitions. Letters above the bars indicate the significant (*p* < 0.05) differences among applied treatments.

### 3.3. Plant Nutrient Concentrations and Recovery

Application of synthetic fertilizer and dairy manure led to distinct pattern of nutrient partitioning in plant parts. Total N, K and S concentrations were significantly different in leaves whereas P concentration was not differed under applied treatments. However, P concentration was significantly different in corn stalks whereas total N, K and S concentrations were similar among applied treatments in corn stalks. Highest leaf N (Figure 5A), K (Figure 5B) and S (Figure 5C) were observed under application of synthetic fertilizer followed by application of manure at 5 Tons. This aligns with synthetic fertilizer response which provide nutrients in readily available forms (Melara et al., 2024) and further indicates that readily available nutrients from synthetic fertilizer were rapidly mobilized to leaves which are primary site for photosynthesis (Cakmak and Engels, 2024). The N, K and S concentration in leaf were similar under higher manure applications and non-fertilized control. Phosphorus concentration in stalk was highest under non-fertilized control followed by higher manure applications at 10 and 15 tons, 5 tons and synthetic fertilizer application (Figure 5D). Nitrogen concentration was affected by the effect of applied treatments, and it was significantly different in corn cobs whereas no significant differences were observed for P, K and S concentrations in cobs (Table 2). Higher N concentrations were found in cobs under non-fertilized control and application of synthetic fertilizer whereas it was statistically similar under manure applications (Figure 5E). These findings suggest that the gradual mineralization of nutrients from higher manure application failed to meet crop demand leading to nutrient deficiency.

**Table 2.** *F*–values and significance obtained through analysis of variance of nutrient concentrations in corn leaf, stalk and cobs and nutrient recovery under different applied treatments.

| Plant Parts | Nutrient Concentration |  |  |  |
| --- | --- | --- | --- | --- |
|  | N | P | K | S |
| Leaves | 20.73*** | 2.63ns | 14.1*** | 8.37** |
| Stalk | 0.94ns | 17.53*** | 0.81ns | 0.58ns |
| Cobs | 3.30* | 0.30ns | 2.58ns | 0.49ns |
| Effects | Nutrient Recovery |  |  |  |
|  | N | P | K | S |
| Treatments | 15.21*** | 2.78ns | 33.25*** | 20.91*** |

\*\*\*: $p < 0.001$ , \*\*: $p < 0.01$ , \*: $p < 0.05$ , ns: non-significant difference at $p \geq 0.05$ .

**Figure 5:**
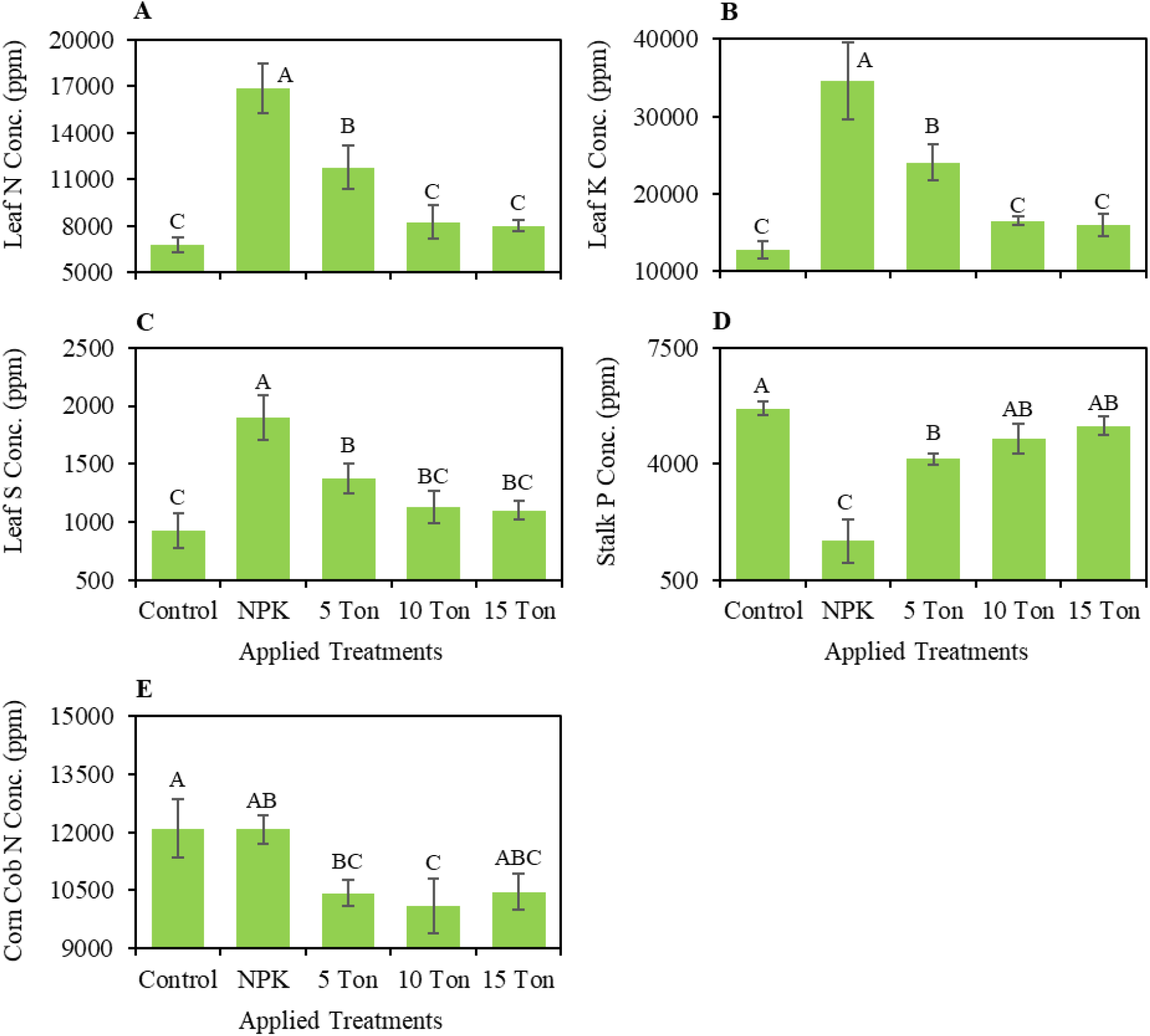
Leaf nitrogen (**A**), potassium (**B**) and sulfur (**C**), stalk phosphorus (**D**) and corn cob nitrogen (**E**) concentration under non-fertilized control, application of synthetic fertilizer and manure application at 5, 10 and 15 tons per acre. Vertical bars indicate ± standard errors for means of four repetitions. Capital letters above the bars indicate the significant (*p* < 0.05) differences among applied treatments.

Nutrient recovery considering N, K and S was significantly different whereas P recovery remained similar under applied treatments (Table 2). Recovery for N, K and S in field corn was highest under application of synthetic fertilizer followed by manure application at 5 tons, 10 and 15 tons and in non-fertilized control (Figure 6). Nitrogen recovery was 259 and 197 lbs acre^-1^ whereas potassium recovery was 438 and 287 lbs acre^-1^ for synthetic fertilizer application and manure application at 5 tons, respectively. Sulfur recovery was 28 and 20 lbs acre^-1^ for synthetic fertilizer application and manure application at 5 tons, respectively (Figure 6). These results also relate to the fact of synthetic fertilizer efficiency to provide readily available nutrients. Similar P response and recovery indicated low P mobility in the given experimental conditions and soils. Nutrient availability in soils and plant uptake cannot be overlooked as various fertilizer types and soil properties impact overall nutrient dynamics (Sistani et al. 2010). While synthetic fertilizers provide readily available nutrients and meet crop nutrient demands, continuous use of these fertilizers impacts soil quality. Hence a balanced approach such as integrating manure application would be beneficial.

**Figure 6:**
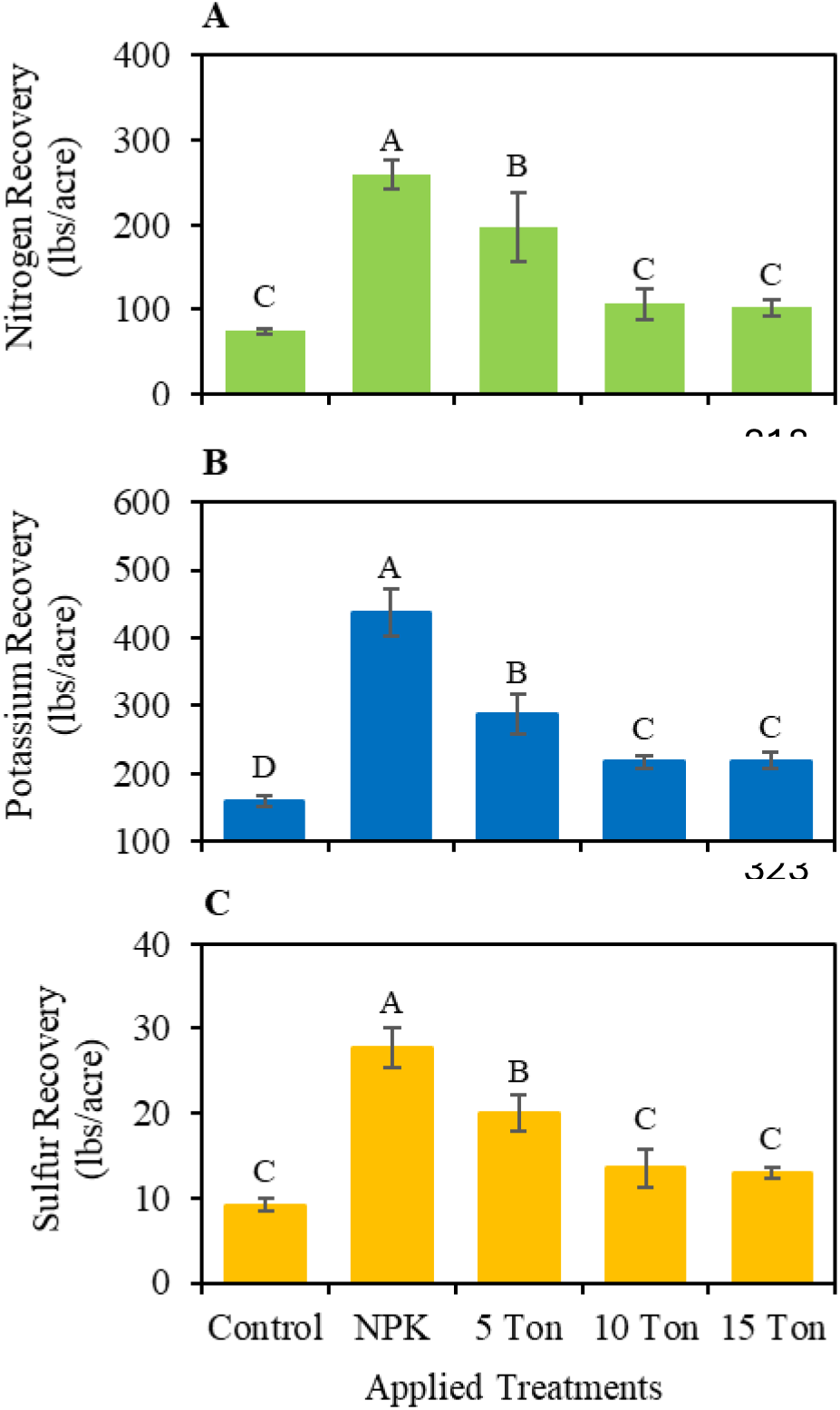
Nitrogen (**A**), potassium (**B**) and sulfur (**C**) recovery under non-fertilized control, application of synthetic fertilizer and manure application at 5, 10 and 15 tons per acre. Vertical bars indicate ± standard errors for means of four repetitions. Capital letters above the bars indicate the significant (*p* < 0.05) differences among applied treatments.

### 3.4 Soil Nutrients

Soil nutrient analysis indicated no significant impact on soil nutrient concentration among treatments except for NH_4_^+^–N (Table 3). Nutrient concentration significantly varied under 0-30, 30-60 and 60-90 cm soil depths. Although non-significant, soil nitrate-N concentration was higher under NPK treatment whereas it was identical under control and manure applications in 0-30 cm soil depths. This indicates that chemical fertilizer application led to higher nitrate-N concentration in soil. Nitrate-N concentrations at 30-60 and 60-90 cm soil depths were similar in all treatments (Figure 7A). Soil NH_4_^+^–N concentration was significantly different under manure applications and depths (Figure 7B). NH_4_^+^–N concentration was higher in upper layer (0-30 cm) whereas it was lower and statistically similar in 30-60 and 60-90 cm soil depths. Under chemical fertilizer and manure applications highest and lowest NH_4_^+^–N was observed in 5 tons and 15 tons of manure application, respectively (Figure 7B). A meta-analysis conducted by Brien and Hatfield (2019) indicated that soil N was not different under synthetic fertilizers and manure applications. The purpose of manure applications is to match the N availability between synthetic fertilizer or manure applications. As soil N pool is dependent upon sample timing and depth, we observed that NO_3_^−^–N and NH_4_^+^–N concentrations were significantly different in upper and lower depths however they were statistically similar but numerically lower in manure application and synthetic fertilizer treatment. Meta analysis conducted by Brien and Hatfield (2019) also suggested that the nitrate leaching may be reduced in manure applications. This may be related to the fact that higher C:N ratio in manure fertilized soils may result in great immobilization and less nitrate availability to losses (Fan et al., 2017). Sampling at the end of season does not account for post-harvest mineralization process, therefore there are higher risks associated with manure applications. This becomes critical if the second crop is not planted to utilize the excessive nutrients (Thapa et al., 2018).

**Table 3.** *F*–values and significance obtained through analysis of variance of nutrient concentrations under different applied treatments and at different soil depths.

| Effects | NO<sub>3</sub><sup>-</sup>-N | NH<sub>4</sub><sup>+</sup>-N | P<sub>2</sub>O<sub>5</sub> | K<sup>+</sup> | SO<sub>4</sub>-S |
| --- | --- | --- | --- | --- | --- |
| Treatment | 2.08ns | 3.68** | 0.83ns | 0.16ns | 1.92ns |
| Depth | 77.25*** | 35.70*** | 31.06*** | 47.10*** | 49.56*** |
| Interaction | 1.85ns | 0.25 ns | 0.70 ns | 1.04 ns | 0.86 ns |
| CV | 41.98 | 25.59 | 50.56 | 24.57 | 27.98 |

\*\*\*: $p < 0.001$ , \*\*: $p < 0.01$ , \*: $p < 0.05$ , ns: non-significant difference at $p \geq 0.05$ .

**Figure 7:**
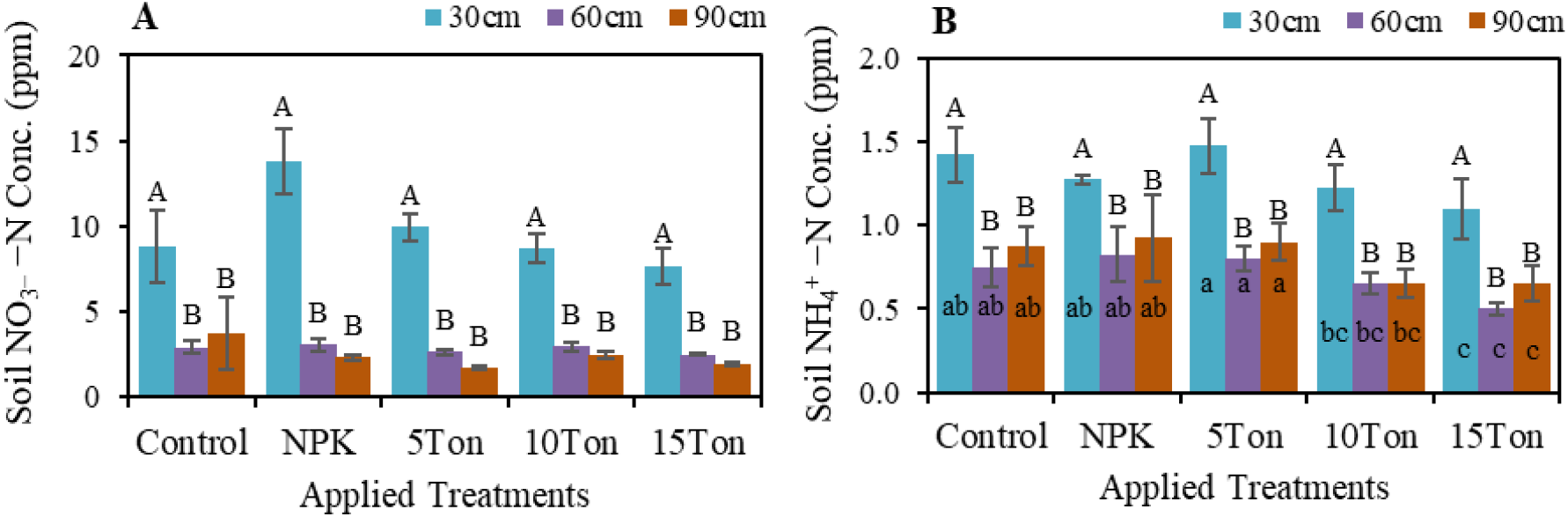
Impact of synthetic fertilizer and manure application rates on soil nitrate-N (**A**) and ammonium-N (**B**) under different soil depths in field corn. Vertical bars indicate ± standard errors for means of four repetitions. Capital letters above the bars indicate the significant (*p* < 0.05) differences among depths within the same treatment whereas small letters inside the bars indicate significant (*p* < 0.05) differences among different applied treatments. Bars without small letters indicate no significant differences.

There were no significant effects for P, K and S concentrations among different treatments; however, concentration of P, K and S was significantly higher in upper soil layer (0-30 cm) and it was lower at 30-60 and 60-90 cm soil depths (Figure 8, 9). Soil K concentration was numerically higher under manure application and significantly different at 30-60 and 60-90 cm soil depths in all treatments (Figure 8B). Soil P (Figure 8A) and S (Figure 9) concentrations were statistically similar at 30-60 and 60-90 cm soil depths. Soil P was statistically similar synthetic fertilizer and manure applications. Although non-significant soil P under 15-ton manure applications were numerically higher compared to 5- and 10-ton manure applications. Increase in soil P under manure application has been reported in previous research (Brien and Hatfield, 2019). Higher soil P is also consistent with low soil P mobility in most soils. Phosphorous also tends to be absorbed by soil particles and organic matter forming insoluble compounds that limit its downward movement (Muniya and Dixit, 2023). These findings on higher soil P are consistent with previous studies (Pan et al. 2024, Anthonio et al., 2023, Wang et al., 2022). Higher concentration of soil K in 0-30 cm soil depth and significant differences between 60 cm and 90 cm soil depths indicate moderate K mobility and redistribution compared to soil P. Higher K in topsoil possibly resulted from direct application of synthetic fertilizer and manure applications. Numerically higher K in soil also suggests that manure could be a greater source for K availability that enriched the soil profile. Sulfur concentration also followed a similar pattern, and higher S was observed at 0-30 cm soil depth. Sulfur exists both in organic and inorganic forms and organic S typically exists in surface layers particularly under manure applications (Zhang et al., 2019). Mineralization of organic matter release sulfate-S which is considered comparatively mobile and can be leached. Statistically similar response of soil S compared to P for 0-30 cm and 30-60 and 60-90 cm soil depths suggest a balance of soil S and its downward mobility. While soil nutrients are influenced by several factors and soil processes, crop performance under different treatment applications may also influence overall plant uptake thus impacting soil nutrient availability. This could be reason for higher soil nutrients where the field corn yield in this was lowest compared to synthetic fertilizer and manure applications.

**Figure 8:**
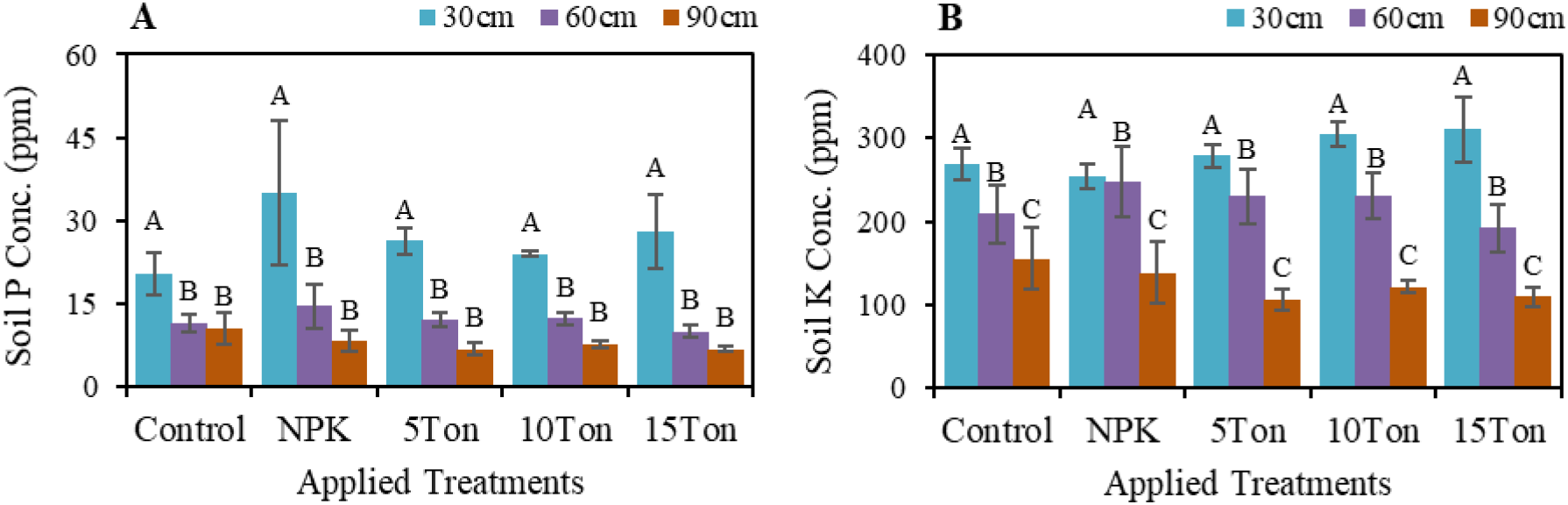
Impact of synthetic fertilizer and manure application rates on soil P (**A**) and K (**B**) under different soil depths in field corn. Vertical bars indicate ± standard errors for means of four repetitions. Capital letters above the bars indicate the significant (*p* < 0.05) differences among depths within the same treatment whereas no significant differences were observed among different applied treatments.

**Figure 9:**
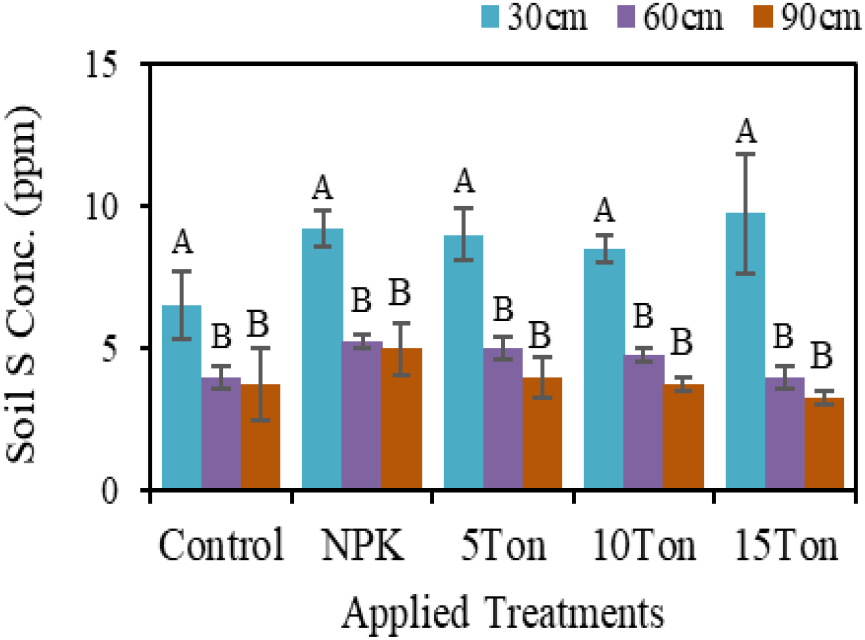
Impact of synthetic fertilizer and manure application rates on soil S under different soil depths in field corn. Vertical bars indicate ± standard errors for means of four repetitions. Capital letters above the bars indicate the significant (*p* < 0.05) differences among depths within the same treatment whereas no significant differences were observed among different applied treatments.

## 4. Conclusions

In-season leaf nutrient concentration was significantly different among applied treatments with the highest concentrations under synthetic fertilizer application. Application of synthetic fertilizer resulted in the highest plant height (116 in) and produced higher corn yield (45.5 tons acre^-1^) compared to non-fertilized control and application of dairy manure. Application of manure at 5 tons acre^-1^ produced higher corn yield (35.5 tons acre^-1^) compared to manure application at 10 (25.9 tons acre^-1^) and 15 tons acre^-1^ (26.1 tons acre^-1^). A similar trend was observed for leaf, stem and cobs fresh and dry weights. Nutrient recovery was higher under application of synthetic fertilizer followed by application of manure at 5 tons acre^-1^. Soil nutrient analysis indicated no significant impact on N, P, K and S concentration among treatments except for NH_4_^+^–N. However, nutrient concentration significantly varied under different soil depths. Although non-significant, soil nitrate-N concentration was higher under application of synthetic fertilizer whereas it was identical under control and manure applications in 0-30 cm soil depths. Results suggest that a combination of synthetic fertilizer application and manure might be a practical approach for balanced nutrient supply for field corn. Lower nutrient recovery and corn yield under excessive manure application indicate a nutrient imbalance or other factors impacting field corn performance. Further investigations are required to explore the potential of manure application to ensure balanced nutrient supply, improved yields and reduced nutrient losses in field corn.

## Author contributions

T.H: Conceptualization; methodology; investigation; data collection, data curation; formal analysis; visualization; writing – original draft; funding acquisition; project management; MFA: contribution in finalizing original draft, writing – review and editing, visualization. Authors agreed to the current version of manuscript.

## Acknowledgement

This research is part of one of the postdoctoral fellowship requirements and associated project responsibilities at Oregon State University. Postdoctoral scholar Tajamul Hussain gratefully acknowledges Oregon State University for providing the platform to carry out this work. The author also thanks temporary staff and HAREC field personnel for their assistance with field operations.

